# Reclassification of seven honey bee symbiont strains as *Bombella apis*

**DOI:** 10.1101/2020.05.06.081802

**Authors:** Eric A. Smith, Kirk E. Anderson, Vanessa Corby-Harris, Quinn S. McFrederick, Audrey J. Parish, Danny W. Rice, Irene L. G. Newton

**Affiliations:** Department of Biology, Indiana University, Bloomington, Indiana, USA; Carl Hayden Bee Research Center, USDA-ARS, Tucson, Arizona, USA; Department of Entomology, University of California, Riverside, California, USA

**Keywords:** Bacterial classification, honey bee, evolution, phylogenetics, average nucleotide identity

## Abstract

Honey bees are important pollinators of many major crops and add billions of dollars annually to the US economy through their services. Recent declines in the health of the honey bee have startled researchers and lay people alike as honey bees are agriculture’s most important pollinator. One factor that may influence colony health is the microbial community. Although honey bee worker guts have a characteristic community of bee-specific microbes, the honey bee queen digestive tracts are colonized predominantly by a single acetic acid bacterium tentatively named “Parasaccharibacter apium”. This bacterium is related to flower-associated microbes such as *Saccharibacter floricola*, and initial phylogenetic analyses placed it as sister to these environmental bacteria. We used a combination of phylogenetic and sequence identity methods to better resolve evolutionary relationships among “P. apium”, strains in the genus *Saccharibacter*, and strains in the closely related genus *Bombella*. Interestingly, measures of genome-wide average nucleotide identity and aligned fraction, coupled with phylogenetic placement, indicate that many strains labeled as “P. apium” and *Saccharibacter* sp. are all the same species as *Bombella apis*. We propose reclassifying these strains as *Bombella apis* and outline the data supporting that classification below.

## Introduction

The honey bee (*Apis mellifera*) is extremely important economically because of the pollination services it provides to numerous agricultural crops. As a result, there is increasing interest in determining how the microbiome supports and influences bee function. While a honey bee colony is made up of individuals with distinct roles, or castes, the majority of studies on bee microbiomes have focused on workers. The microbial community of worker bees consists of eight to ten core bacterial phylotypes (1, 2). The characterization of these groups led to speculation about their role in honey bee health and whether or not they provision nutrients (1) or assist in the breakdown of plant-derived carbohydrates (3, 4), as is the case in other insect-microbe interactions (5, 6). There has also been speculation as to the role of the microbiome in resistance to pathogens, as microbial communities have been shown to protect the bumble bee (*Bombus terristris*) from the parasite *Crithidia bombi* (7). Honey bee-associated microbes interact with each other in diverse ways both *in vitro* and *in vivo*, suggesting that they may interact syntrophically within workers (8, 9). While these studies focused on honey bee workers are intriguing, it is surprising that only recently was the microbiome of queen bees throughout development characterized (10).

Interestingly, the microbial community associated with queen bees is vastly different than that of workers and comprises a large percentage of acetic acid bacteria, a group of bacteria present only at very small percentages in workers. One of the primary bacteria that differentiate queens from workers was tentatively named “Parasaccharibacter apium” (11). This bacterium is in the family *Acetobacteraceae* and occupies defined niches within the hive, including: queen guts, nurse hypopharyngeal glands, nurse crops, and royal jelly, and is only rarely found in high abundance outside of these areas (12, 13). Evidence suggests that it might play a role in protecting developing larvae and worker bees from fungal pathogens, such as *Nosema* (11, 14, 15). Given that this bacterium makes up a large proportion of the queen gut microbiome, it is possible that it plays an important role in queen nutrition, protection from pathogens, and possibly modulating queen fertility, fecundity, and longevity (16).

We sought to determine the evolutionary relationships between strains of “P. apium”, strains of the closely related genus *Saccharibacter*, and characterized sequences from strains of the recently named genus *Bombella* (17), using previously published data (see Supplementary Table 1 for information on all genomes used in this study). Using a combination of phylogenetic and sequence similarity methods, we found that many genomes labeled as “P. apium” or *Saccharibacter* sp. are actually the same species as strains of the previously described species *Bombella apis*. We reclassify these strains and outline the data supporting this reclassification below.

## Materials and methods

### 16S rRNA phylogeny

To determine relatedness of the strains classified as “P. apium*”* and *Saccharibacter* spp. compared to existing *Bombella* and *Saccharibacter* species, we compared 16S rRNA gene sequences of these genomes to one another. We first downloaded all 16S rRNA gene sequences from NCBI that were labeled as *Bombella*, “Parasaccharibacter”, or *Saccharibacter* (Supplementary Table 1). We also included *Gluconobacter oxydans* DSM3503 for use as an outgroup. All sequences were aligned using the SINA aligner (Pruesse *et al*., 2012); parameters used were set using the --auto option. A maximum likelihood phylogeny was constructed using RAxML with the GTRGAMMA substitution model and 1000 bootstrap replicates (v8.2.11, (18)). The final tree was visualized using ggtree in R (v3.12 https://bioconductor.org/packages/release/bioc/html/ggtree.html).

### Sequencing and assembly of Bombella apis MRM1

*Bombella apis* MRM1 was requested of the Riken JCM repository (31623^T^) and the microbe streaked to isolation on de Man, Rogosa, Sharpe (MRS) agar at 34C. A single isolated colony was used to inoculate MRS broth and grown overnight at 34C, 250 rpm before harvesting cells for DNA extraction. DNA was isolated using a standard beat beating method followed by phenol:chloroform extraction. Libraries were generated using an NEBNext Ultra II DNA sequencing library kit and sequenced on a NextSeq instrument using 300 cycles and a paired-end sequencing strategy. Reads were binned by their barcode and Masurca was used to assemble the reads using the guided assembly approach with the *Bombella apis* A29 genome as a reference. The resulting assembly was quality controlled using Quast and CheckM and comprises 6 contigs, with an N50 of 411,263 and a total size of 2,048,899 bp. This Whole Genome Shotgun project has been deposited at DDBJ/ENA/GenBank under the accession JAGJTM000000000. The version described in this paper is version JAGJTM010000000.

### Core ortholog phylogeny

To determine genome-wide phylogenetic relationships between strains, we clustered genes from all genomes in Supplementary Table 1 into orthologous gene clusters using reciprocal blast. Whole genome and amino acid sequences were downloaded from NCBI (see accessions in Supplementary Table 1) and clustering was performed using default parameters and all vs. all BLASTp with default options. An orthologous group of genes was defined by complete linkage such that all members of the group had to be the reciprocal best hit of all other members of the group. Thus, a particular strain could have at most one gene per ortholog group. We constructed a phylogeny using concatenated amino acid alignments of all single-copy orthologs. The amino acid sequences were aligned using the MAFFT L-INS-I algorithm (v7.310, (Katoh *et al*., 2002)), and alignments were then concatenated, and used to construct a maximum likelihood phylogeny using RAxML with substitution model GTRGAMMA model and 1000 bootstrap replicates (v8.2.11, (Stamatakis, 2006)). The final tree was visualized using ggtree in R (v3.12 https://bioconductor.org/packages/release/bioc/html/ggtree.html).

### Calculation of genomic similarity

To determine relatedness and proper species assignment, we calculated genome-wide Average Nucleotide Identity (gANI) and aligned fraction (AF) for each pairwise comparison using ANIcalculator (Varghese *et al*., 2015). Predicted transcript sequences for each pairwise comparison were passed to the software, which output gANI and AF in each direction for the pairwise comparison. As gANI and AF can vary depending on the direction of comparison due to differences in genome lengths, we report results for comparisons in both directions.

## Results

### 16S rRNA gene phylogeny

To determine phylogenetic relationships of all genomes labeled “Parasaccharibacter”, *Saccharibacter*, and *Bombella* on NCBI, we constructed a maximum likelihood phylogeny using 16S rRNA sequences (Figure 1A). This phylogeny, based on the 16S rRNA gene sequences from the following genomes, suggests that the strains are all very closely related and represent a single, distinct clade: *Saccharibacter* sp. AM169, “Parasaccharibacter apium” G7_7_3c, “Parasaccharibacter apium” A29, “Parasaccharibacter apium” B8, “Parasaccharibacter apium” C6, *Sacchahribacter* sp. M18, *Saccharibacter* sp. 3.A.1, *Bombella apis* MRM1, *Bombella apis* SME1. For simplicity, this clade, highlighted in yellow in Figure 1, will hereafter be referred to as the “clade of interest”. “P. apium” AS1 is not included in the clade of interest for reasons outlined below. The relationships between strains within the clade of interest are difficult to determine – as is evidenced by the low bootstrap support for nodes within the clade – owing to the high degree of similarity between the 16S rRNA gene sequences (Supplementary Table 2). However, the phylogeny clearly places these accessions with *B. apis*, and sister to “P. apium” AS1. It is interesting to note that, although the sequences all show similar degrees of divergence from *B. intestini* and *B. apis*, bootstrap support is quite high for the separation of *B. intestini* from the clade of interest (Figure 1A).

**Figure 1.**
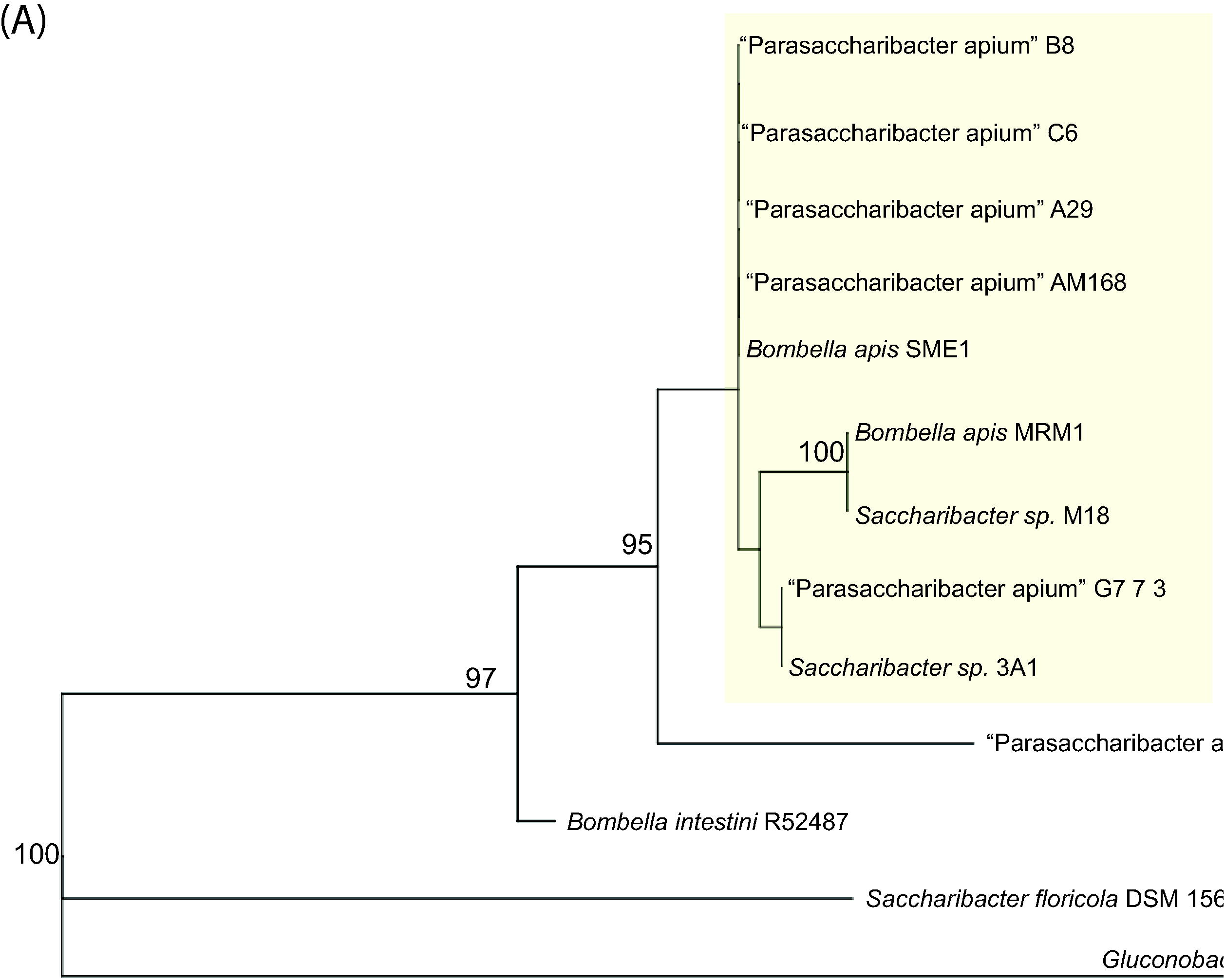
Maximum likelihood 16S rRNA gene phylogeny (A) and core ortholog phylogeny (B). Numbers at nodes represent bootstrap support from 1000 bootstrap pseudoreplicates; only support greater than 80 shown. The “clade of interest” is highlighted in yellow.

### Core ortholog phylogeny

We reciprocal blast to define core orthologs using the “Parasaccharibacter”, *Saccharibacter*, and *Bombella* genomes listed in Supplementary File 1; *Gluconobacter oxydans* DSM3503 was used as an outgroup. In total, an alignment of 2,270,622 nucleotide positions was defined, with nucleotide positions represented by 4 or more taxa. This alignment was then used to construct a core ortholog phylogeny (Figure 1B) to better resolve phylogenetic relationships.

This core orthology phylogeny largely agrees with our 16S rRNA gene phylogeny with one exception: the relationships of *B. intestini* and “P. apium” AS1 have switched such that *B. intestini* is now sister to the clade of interest. Bootstrap support for the nodes in this tree are much higher than in the 16S rRNA gene tree, including within the clade of interest. Importantly, bootstrap support among the nodes making up the main backbone of the tree are all 100, indicating very high support for the basal nodes (Figure 1B).

### Calculation of genomic similarity

Given the discrepancy between nomenclature and the phylogeny (the placement of “Parasaccharibacter” and *Saccharibacter* spp. within the *Bombella* group), and considering the short branch lengths within the clade of interest, we calculated genome-wide Average Nucleotide Identity (gANI) and aligned fraction (AF) to clarify species relationships (Figure 2). The general criteria for two genomes to be considered the same species are AF > 0.6 and gANI > 96.5 (19).

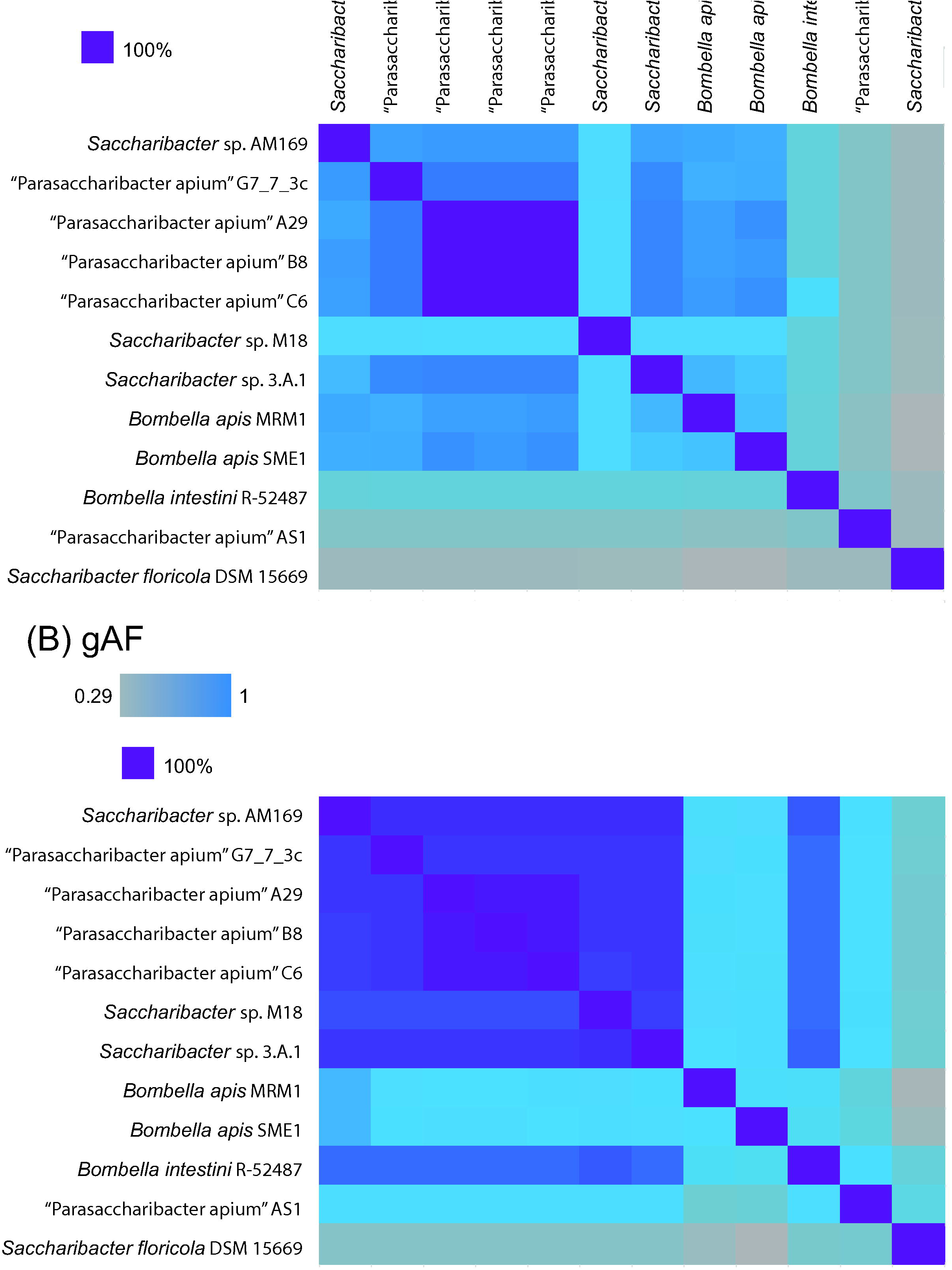

Using these criteria, all genomes within the clade of interest should be considered the same species, and distinct from *B. intestini* and “P. apium” AS1 (Supplementary Table 3), though part of the *Bombella* genus. Both the high degree of support for placement of *B. apis* within the clade of interest and the gAF and gANI results strongly suggests that the genomes within the clade of interest belong to the same species and are all *Bombella apis* strains.

## Discussion

Here, we used a combination of phylogenetic analysis (16S rRNA gene and core orthologs) and calculations of aligned fraction and genome-wide average nucleotide identity to determine relationships among symbionts of honey bees, bumble bees, and environmental bacteria. The phylogenetic data largely agree with each other, and with the AF and gANI delimitations of species. The combination of these data indicate that the genomic accessions GCA_000723565.1, GCA_002079945.1, GCA_002917995.1, GCA_002917945.1, GCA_002917985.1, GCA_002150105.1, and GCA_002150125.1 are all the same species, despite being named in NCBI as various strains of “Parasaccharibacter apium” and *Saccharibacter* sp. Given that the species name “P. apium” has been effectively but not validly published (i.e. in accordance with IJSEM standards)(Corby-Harris *et al*., 2016), while *B. apis* has been validly published (Yun *et al*., 2017), we propose renaming the above accession numbers to reflect the proper genus and species assignment while maintaining their current strain designations (see Table 1 for full genus, species, and strain designations).

**Table 1.** GenBank accession number, current name, isolation source, genome size, %GC, and proposed new names of the strains used in this study.

| GenBank accession | Current name | Isolation source | References | Genome size (Mbp) | %GC | Propose |
| --- | --- | --- | --- | --- | --- | --- |
| GCA_002079945.1 | “Parasaccharibacter apium” G7_7_3c | *Apis mellifera* hindgut | Corby-Harris & Anderson 2018 | 2.01 | 59.42 | *Bombel* |
| GCA_002917995.1 | “Parasaccharibacter apium” A29 | *Apis mellifera* larva | Corby-Harris & Anderson 2018 | 2.01 | 59.39 | *Bombel* |
| GCA_002917945.1 | “Parasaccharibacter apium” B8 | *Apis mellifera* larva | Corby-Harris & Anderson 2018 | 2.01 | 59.38 | *Bombel* |
| GCA_002917985.1 | “Parasaccharibacter apium” C6 | *Apis mellifera* larva | Corby-Harris & Anderson 2018 | 2.01 | 59.38 | *Bombel* |
| GCA_000723565.1 | “Parasaccharibacter apium” AM168 | *Apis mellifera* stomach | Chouaia et al. 2014 | 1.98 | 59.32 | *Bombel* |
| GCA_002150105.1 | *Saccharibacter* sp. M18 | *Apis mellifera* stomach | Veress et al. 2017 | 2.01 | 59.35 | *Bombel* |
| GCA_002150125.1 | *Saccharibacter* sp. 3.A.1 | Honey | Veress et al. 2017 | 2.01 | 59.41 | *Bombel* |
| GCA_009362775.1 | *Bombella apis* SME1 | Honey | Smith et al., 2019 | 2.09 | 59.56 | N/A |
| PRJNA719376 | *Bombella apis* MRM1 | *Apis mellifera* midgut | Yun et al., 2017 | 2.01 | 59.39 | N/A |

## Authors and contributors

Conceptualization: EAS and ILGN; Sequencing and assembly: AJP, Formal analysis: EAS, DWR; Writing – original draft: EAS; Writing – review and editing: EAS, KEA, VCH, QSM, ILGN

## Conflicts of interest

The authors declare that there are no conflicts of interest.

## Funding information

EAS is supported by Agriculture and Food Research Initiative -Education and Workforce Development project accession no. 1019114 from the USDA National Institute of Food and Agriculture

## Acknowledgements

The authors thank Amelia R. I. Lindsey, Delaney L. Miller, and Audrey J. Parish for helpful feedback on drafts of this manuscript.

---

**Supplementary Table 1**. GenBank accession numbers for all genomes used in this study.

**Supplementary Table 2**. Pairwise percent identity of 16S rRNA gene sequences for the genomes of the clade of interest. Colors in each cell scale with the metric from 0-100.

**Supplementary Table 3**. Pairwise aligned fraction and genome-wide average nucleotide identity of all strains used in this study. Colors in each cell scale with the metric. Aligned fraction scales from 0-100, while gANI scales from 0-1. To be considered the same species, two genomes should have aligned fraction > 0.6 and gANI > 96.5 (19). Both metrics can vary depending on which genome is considered the reference for alignment, and we made the calculations in both directions. Genomes listed on the horizontal axis were considered as the reference for these calculations.

## Notes

### Competing Interest Statement

The authors have declared no competing interest.

### Summary of Updates

This manuscript has been revised to include the genomic comparisons with the type strain, Bombella apis MRM1

